# Modeling the Impact of Contamination on Microbial-Enzyme Dynamics and Soil Function

**DOI:** 10.64898/2026.10.07.757410

**Authors:** Sepideh Vafaie, Nina M. Goodey, Jennifer Adams Krumins, Eric Forgoston

## Abstract

Globally, contaminated soils pose serious risks to agriculture, biodiversity of ecosystems, and human health. Although a variety of monitoring and remediation methods have been developed over the years, there remains a lack of theoretical modelling to better understand the impact of contamination on soil carbon in a microbially-mediated system. In this study, we develop a model which incorporates contaminated soil organic carbon, uncontaminated soil organic carbon, dissolved organic carbon, microbes, extracellular enzymes, and CO_2_, to explore how different amounts of contamination and modes of contamination (pulse and press) affect microbially-mediated carbon flux within both vegetated and barren environments. The model represents contamination as a pool of contaminated soil organic carbon that is more resistant to microbial degradation than uncontaminated soil organic carbon. The results show that, in barren environments, the amount and type of contamination has no notable impact on the soil system. In contrast to this, in vegetated environments, plants mitigate the impact of large contamination inputs with differences in outcome depending on the amount and type of contamination.

**Highlights:**

- Developed a microbial-enzyme model to investigate contaminated soil carbon dynamics.
- Studied pulse and press contamination within vegetated and barren environments.
- In barren environments, contamination has no notable impact on the soil system.
- Vegetation mitigates the impact of large contamination inputs.
- Pulse contamination results in slightly better long-term recovery than press contamination.

## 1. Introduction

Soil microbial biomass maintains soil health by driving carbon and nutrient cycling, which supports ecosystem sustainability (Singh and Gupta, 2018). Enzyme activity is a key indicator of soil health, with most enzymes originating from soil microorganisms and some from plants and animals (Nannipieri, Kandeler and Ruggiero, 2002). Soil microbes interact with soil organic carbon by producing extracellular enzymes that break down complex organic molecules and release nutrients (Kelley, Fay, Polley, Gill and Jackson, 2011). The presence of diverse plant species further enhances microbial activity, promoting efficient nutrient cycling, particularly nitrogen cycling (Zak, Holmes, White, Peacock and Tilman, 2003). Soil-emitted CO_2_ and dissolved organic matter (DOM), a mobile and reactive form of carbon, both play important roles in global carbon cycling (Elberling and Brandt, 2003).

Soil contamination creates serious risks to agriculture (Hou, O’Connor, Igalavithana, Alessi, Luo, Tsang, Sparks,Yamauchi, Rinklebe and Ok, 2020; Alengebawy, Abdelkhalek, Qureshi and Wang, 2021; Rashmi, Roy, Kartika, Pal, Coumar, Kala and Shinoji, 2020; Lu, Song, Wang, Liu, Meng, Sweetman, Jenkins, Ferrier, Li, Luo and Wang, 2015), biodiversity (Wang, Teng, Wang, Xu, Li, Sun, Hu, Zhao, Ren and Luo, 2022; van Straalen, 2002; Raimi, Iyingiala, Sawyerr, Saliu, Ebuete, Emberru, Sanchez and Osungbemiro, 2022), human health (Ghosh, Manna, Jha, Singh, Misra, Srivastava, Srivastava, Laik, Bhattacharyya, Prasad, Singh, Singh, Kumar, Tiwari and Singh, 2022; Steffan, Brevik, Burgess and Cerdà, 2018; Brevik and Sauer, 2015), and water resources (Tripathi, Husain, Ahmad, Hasan and Farooqui, 2021; Ashraf, Maah and Yusoff, 2014; Dehkordi, Nodeh, Dehkordi, Khorjestan and Ghaffarzadeh, 2024). Research has found complex relationships among contamination, microbial communities, extracellular enzyme activity, and soil carbon cycling, with responses varying by contaminant type, concentration, vegetation, and soil properties (Hagmann, Goodey, Mathieu, Evans, Aronson, Gallagher and Krumins, 2015). For example, studies of contaminated soils have shown that heavy metals alter soil microbial community composition, enzyme activity, and soil carbon dynamics (Singh, Vaidya, Goodey and Krumins, 2019b). Additional studies have shown that contaminated soils often contain more persistent organic matter and that abiotic soil properties can have a greater influence on microbial function than microbial inoculation alone (Semple, Morriss and Paton, 2003; Delgado-Baquerizo, Grinyer, Reich and Singh, 2016; Hagmann, Kruge, Cheung, Mastalerz, Gallego, Singh, Krumins, Li and Goodey, 2019; Singh, Ojinnaka, Krumins and Goodey, 2019a; Zeng, Huang, Guo, Dai, He, Chen and Xin, 2024).

A profound challenge for environmental engineers is to resolve the mechanisms of microbial degradation of organic contaminants (Li, Cai, Deng, Zhang, Li, Luo and Zhang, 2025). Microbial-mediated decomposition of organic contaminants occurs in a heterogeneous soil matrix containing both contaminated and uncontaminated soil organic carbon. The balance between these carbon pools likely influences microbial metabolism and extracellular enzyme production. To this end, extensive research has been conducted to understand and mitigate soil contamination, particularly with a focus on monitoring and remediation. Various tools, such as terrestrial model ecosystems (TMEs) (van Straalen, 2002) and bioinformatics (van Straalen, 2002; Gao, Wu, Liu, Ou, Zhang and Duan, 2023), are used to assess the impact of soil contamination on ecosystems. Additional studies have explored plant–soil feedback mechanisms as a framework for understanding the ecological dynamics of metal-contaminated soils (Krumins, Goodey and Gallagher, 2015). Research has also shown that organic amendments, including plant derived carbon inputs, can improve soil moisture, organic matter content, and microbial activity in contaminated soils (Liu, Hou, Wang, Yang, Luo and Christie, 2015; Hagmann et al., 2015; Vaidya, Hagmann, Haramuniz, Krumins and Goodey, 2022). Additional studies have shown that vegetation and plant derived carbon inputs can enhance microbial activity and soil function, while microbial inoculation alone is often insufficient to revitalize contaminated soils (Lu, Sun and Zhu, 2017; Zhou and Staver, 2019; Vaidya, Hagmann, Balacco, Passchier, Krumins and Goodey, 2020; Vaidya, Krisak, Krumins and Goodey, 2024)

Heavy metal removal strategies include physical, chemical, and combined remediation methods (Khalid, Shahid, Niazi, Murtaza, Bibi and Dumat, 2017), including identification of pollution sources (Shan, Tysklind, Hao, Ouyang, Chen and Lin, 2013; Facchinelli, Sacchi and Mallen, 2001), reduction of metal bioavailability (Zhang and Pu, 2011), and application of bioremediation techniques (Zhao, Ma, Zhu, Tang and McGrath, 2015). In particular, phytomanagement has been effective in removing copper (Cu) from soil, demonstrating success in both short- and long-term applications (Burges, Fievet, Oustriere, Epelde, Garbisu, Becerril and Mench, 2020). Notably, approximately 500 plant species have been identified for phytoremediation, and their ability to absorb heavy metals from contaminated soil has been demonstrated (Li, Zhou, Qin, Tian, Qi, Yan and Han, 2019). In addition to phytomanagement, other methods for cleaning up polluted areas include surface capping (Zhang, Zhu, Zeng, Yu, Cui, Yang and Shen, 2016; Knox, Paller and Roberts, 2012) and soil washing (Liu, Li, Song and Guo, 2018).

However, chemical remediation techniques, including immobilization (Nejad, Jung and Kim, 2018), soil washing (Liu et al., 2018), and vitrification (Sanito, Bernuy-Zumaeta, You and Wang, 2022), pose environmental risks, such as toxic byproducts, soil degradation, and groundwater contamination, limiting their large-scale applications (Murtaza, Murtaza, Niazi and Sabir, 2014). Additionally, because microbial metabolism of contaminated, and potentially toxic, soil carbon occurs on a timescale of decades or centuries (Jensen, Genereux, Solomon, Knappe and Gilmore, 2024), the implementation of rigorous experiments over realistic timelines is essentially impossible. Therefore, mathematical models are required to fully understand these phenomena (Hart, Kulakova, Allen, Simpson, Oppenheimer, Masoom, Courtier-Murias, Soong, Kulakov, Flanagan, Murphy and Kelleher, 2013; Moreno, Poznyak, Rodríguez, Chairez and Dorantes-Rosales, 2020; Raajaraam and Raman, 2024). Several theoretical models have been developed to study soil functioning, but these models focus primarily on uncontaminated soil systems.

Parton et al. (Parton, Schimel, Cole and Ojima, 1987) developed a model to study how climate and soil texture influence soil organic carbon and nitrogen levels in the U.S. Great Plains. This model classifies soil organic matter (SOM) into three categories based on decomposition rates: active SOM (rapid turnover), slow SOM (moderately stable), and passive SOM (highly stable, persisting for over 200 years). By incorporating these factors, the model successfully estimated organic matter levels across different grassland sites in the U.S. (Parton et al., 1987) Molina et al. (Molina, Hadas and Clapp, 1990) developed the NCSOIL (Nitrogen and Carbon Soil model) model to explore two competing theories on nitrogen and carbon cycling in soil: the direct hypothesis and the MIT (mineralization-immobilization turnover) hypothesis. The direct hypothesis suggests that soil microbes assimilate nitrogen directly from organic matter, while the MIT hypothesis proposes that microbes can only absorb nitrogen from organic matter after its conversion to inorganic form (Molina et al., 1990). They found that while both mechanisms may occur simultaneously in soil, empirical data more closely align with the MIT hypothesis (Molina et al., 1990).

Chertov and Komarov (Chertov and Komarov, 1997) developed SOMM (Soil Organic Matter Mineralization), a model that simulates plant litter decomposition, nutrient storage, and nitrogen release in soil. The model incorporates factors such as temperature, moisture, and the ash content of fallen leaves to model soil organic matter dynamics (Chertov and Komarov, 1997). Schimel and Weintraub (Schimel and Weintraub, 2003) studied two models. The first is a carbon-only (C-only) model, which includes microbial biomass, dissolved organic carbon, soil organic matter, and exoenzymes. The second model is a carbon and nitrogen (C and N) model, which additionally accounts for the inorganic contributions of nitrogen and microbial cell biomass within soil organic matter. Their results suggest that carbon decomposition rates in soil are influenced by both carbon availability and enzyme activity. Their results also predicted that increases in soil nitrogen availability decreases respiration rate but speeds up the growth of microbial biomass (Schimel and Weintraub, 2003). Allison et al. (Allison, Wallenstein and Bradford, 2010) developed a model that includes microbial biomass, dissolved organic carbon, soil organic carbon, and enzymes to explore why CO_2_ emissions initially increase with rising temperatures. After the initial increase in CO_2_ emissions following rising temperature, over time the CO_2_ emissions decline due to microbes’ reduced ability to function at high temperatures. Eventually, the CO_2_ emissions rise again as microbes adapt to the warmer conditions (Allison et al., 2010).

Despite numerous experimental studies on contamination and remediation, theoretical models addressing contaminated soil dynamics remain limited. These models are crucial for understanding the long-term impacts of contamination on soil systems and for identifying the most effective strategies for remediating the soil with minimal harm to the environment and human health. This study introduces a novel microbial-enzyme soil model to assess long-term soil dynamics under contamination. Rather than modeling individual contaminants, this model represents their aggregate effect on soil microbial carbon processing. The model distinguishes between press and pulse contamination. Press contamination refers to continuous pollutant input over time, such as a continuous trickle of contaminated wastewater from a factory, while pulse contamination involves a short-term episode of pollutant exposure akin to a one time chemical spill. These distinctions help clarify how microbial and enzymatic processes respond under different contamination regimes. Findings from the model provide insights into the long-term effects of contamination, aiding policymakers and environmental managers in developing effective response strategies.

## 2. Materials and Methods

Our contaminated soil model is based on the framework proposed by Schimel and Weintraub (Schimel and Weintraub, 2003), and later used by Allison et al. (Allison et al., 2010). Different from their implementations, we incorporate contamination into the microbial-enzyme soil model to study its long-term impact on soil systems. In our model, shown in Figure 1, contaminated soil organic carbon is represented as a separate component to assess the impact of contamination on various soil elements.

**Figure 1.**
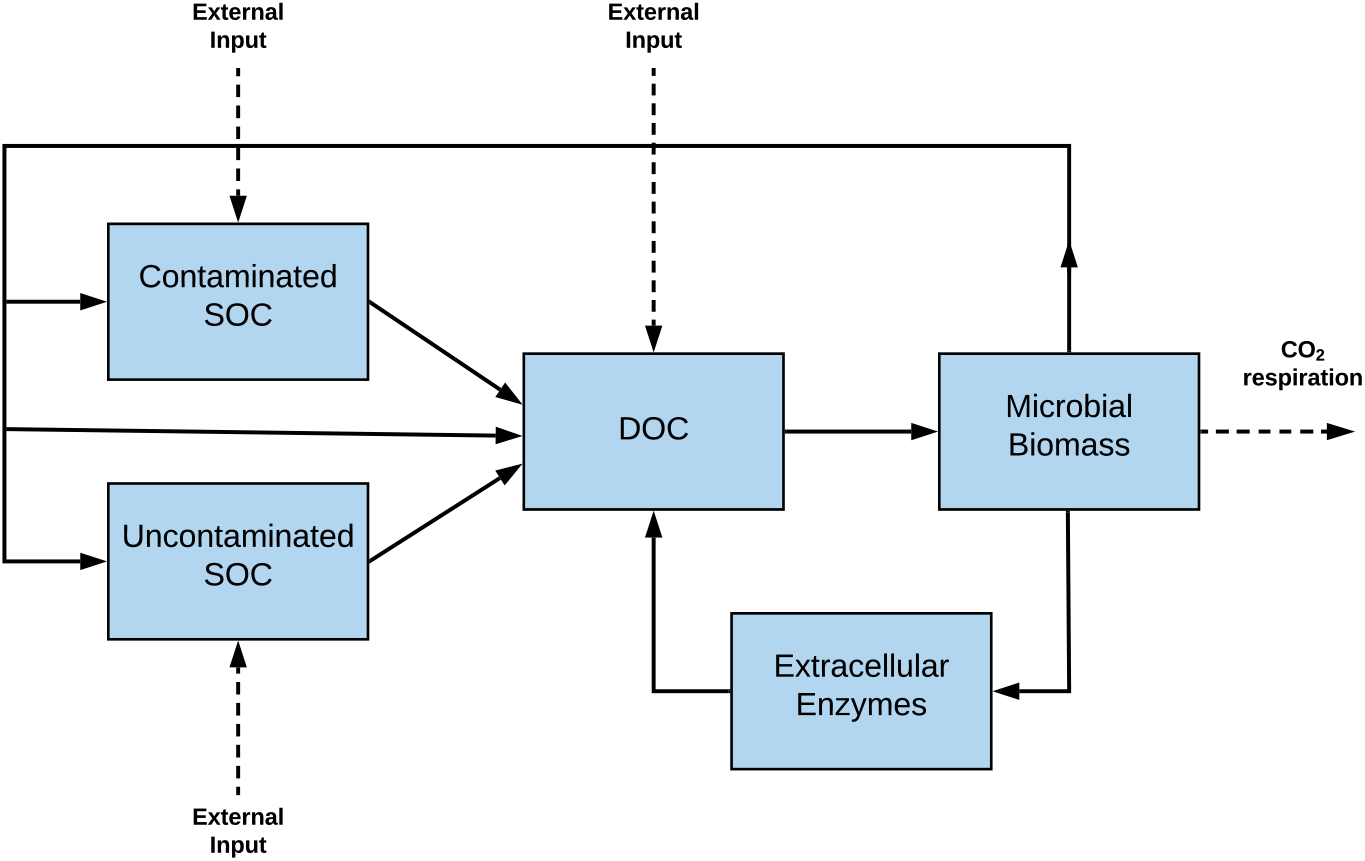
Compartment model showing the decomposition of soil organic carbon in a microbial-enzyme system. Microbial biomass controls the production of extracellular enzymes, which directly affects the conversion of contaminated and uncontaminated soil organic carbon to dissolved organic carbon. By adjusting the parameters, the model can be used to study both vegetated and barren soil environments.

In this model, extracellular enzymes act as a catalyst for the conversion of contaminated soil organic carbon (CSOC) and uncontaminated soil organic carbon (USOC) to dissolved organic carbon (DOC). An important feature of the model is that microbial biomass controls the production of extracellular enzymes, thereby affecting the conversion of CSOC and USOC to DOC. Importantly, we presume that these conversions provide the rate-limiting step in CSOC and USOC decomposition.

Temperature sensitivity has been incorporated into the model as it is well-known that temperature can affect both microbial physiology and extracellular enzyme activity (Davidson and Janssens, 2006; Bradford, Davies, Frey, Maddox, Melillo, Mohan, Reynolds, Treseder and Wallenstein, 2008). Temperature sensitivity of the extracellular enzyme activity is represented via the Arrhenius relationship, (Davidson and Janssens, 2006) while sensitivity of microbial physiology to temperature is represented via carbon use efficiency (CUE). As CUE declines with increasing temperatures, the allocation of carbon to microbial growth is reduced with an accompanying increase in respiration.

By integrating contamination into this framework, we are able to examine how pollutants influence microbial biomass, enzymatic function, and carbon cycling over long periods of time. The approach provides insight into the persistence of soil carbon under environmental stress and informs potential remediation strategies. In the following section, we derive a system of ordinary differential equations (ODEs) which describe how CSOC, USOC, DOC, microbial biomass, and extracellular enzymes change over time in the presence of contamination for both vegetated and barren soil environments.

### 2.1. Model Equation Derivations

Microbial biomass increases due to assimilation of DOC times the carbon use (microbial growth) efficiency, and decreases due to microbial biomass death and enzyme production. The ODE describing the change of microbial biomass in time is given as

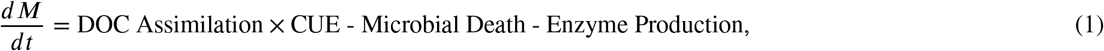

where *M* denotes the microbial biomass, and CUE represents the carbon use efficiency. DOC assimilation follows a Michaelis-Menten equation which has been scaled to the size of *M*, the microbial biomass, so that

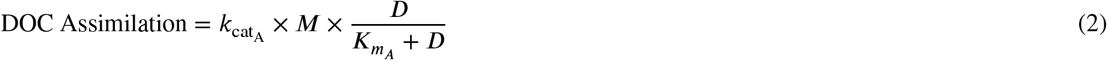

where *D* denotes the dissolved organic carbon, DOC, 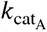, the catalytic constant for assimilation, determines the speed at which the enzyme can catalyze the reaction, and 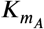represents the Michaelis constant for assimilation. The available cell surface area for uptake is assumed to be directly proportional to the number of cells. Also, microbes are assimilation limited in the sense that they can assimilate no more than the amount of available DOC (Allison et al., 2010).

In Eq. (2), microbial death is proportional to the microbial biomass and is described by

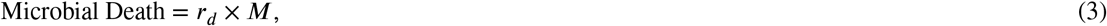

where *r*_*d*_ is a constant death rate. Similarly, enzyme production is also proportional to the microbial biomass, and is described by

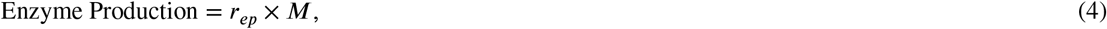

where *r*_*ep*_ is the rate of enzyme production. In contrast to the death rate, the enzyme production rate varies as a linear function of the amount of contaminated and uncontaminated soil organic carbon. Specifically, an increase in the ratio of CSOC to USOC leads to a decrease in the enzyme production rate so that

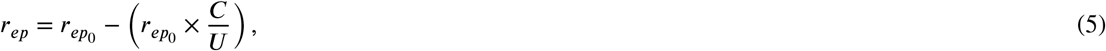

where 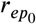is the enzyme production rate in the absence of contamination, and *C* and *U* respectively denote the contaminated soil organic carbon, CSOC, and uncontaminated soil organic carbon, USOC. It is possible for CSOC to increase while USOC remains relatively unchanged. In this scenario, after some amount of time has elapsed, the CSOC to USOC ratio may satisfy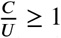. From Eq. (5), this implies that *r*_*ep*_ ≤ 0. If this occurs, we stop the simulation as soon as *r*_*ep*_ crosses the zero thre^*U*^shold.

Substitution of Eqs.(2)-(5) into Eq. (1) gives the final form of the ODE for microbial biomass as

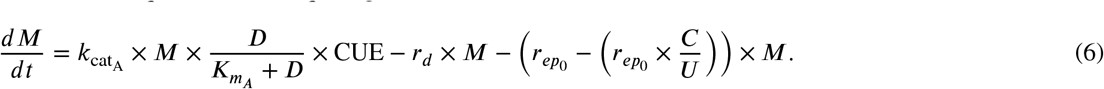

The parameters 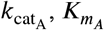, and CUE are all temperature dependent. Using the Arrhenius equation, one can write,

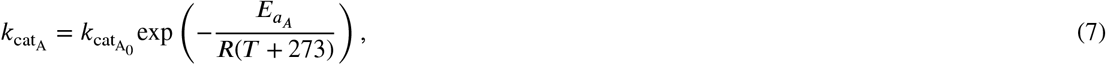

where 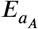is the activation energy needed to convert substrate into product, *R* is the ideal gas constant, *T* is the temperature in degrees Celsius, and 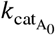 is the pre-exponential coefficient. The parameters *K*_mA_ and CUE are linear functions of temperature so that

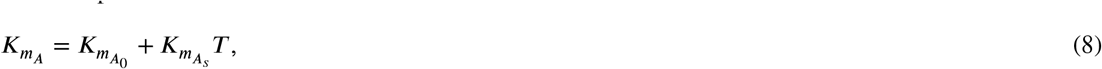

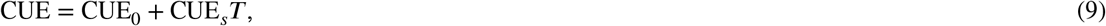

where 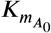and CUE_0_ are respectively the values of 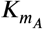 and CUE at 0° Celsius, and 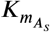and CUE respectively denote the slopes of the linear functions of 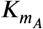and CUE.

Equations (1) and (6) show that only a fraction, CUE, of DOC assimilation contributes to microbial growth. The remaining fraction, 1-CUE, of DOC assimilation is associated with CO_2_ production so that

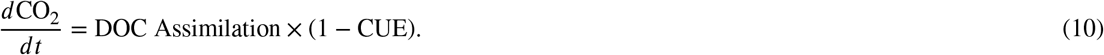

The Extracellular enzymes increase due to enzyme production and decrease with enzyme turnover/loss. The ODE describing the change of extracellular enzymes in time is given as

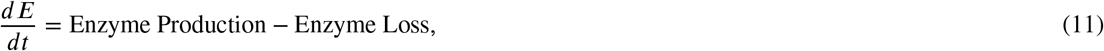

where *E* denotes the extracellular enzymes. The equations for enzyme production, as explained above, are given by Eqs. (4)-(5).

In Eq. (11), enzyme loss is proportional to the extracellular enzymes and is described by

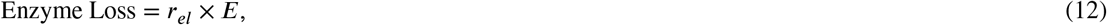

where *r*_*el*_ is a constant enzyme loss rate.

Substitution of Eqs. (4)-(5) and Eq. (12) into Eq. (11) gives the final form of the ODE for extracellular enzymes as

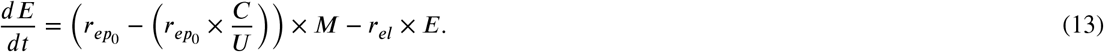

Uncontaminated soil organic carbon, USOC, increases due to external inputs as well as contributions from a fraction of dead microbial biomass, and decreases due to losses from USOC decomposition. The ODE describing the change of USOC in time is given as

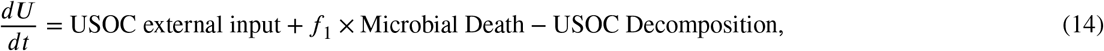

where *U*, as above, denotes the uncontaminated soil organic carbon, USOC, and *f*_1_ represents the fraction of dead microbial biomass which is added to the USOC. The equation for microbial death, as explained above, is given by Eq.(3).

USOC decomposition is catalysed via extracellular enzymes and is modeled with Michaelis-Menten kinetics so that

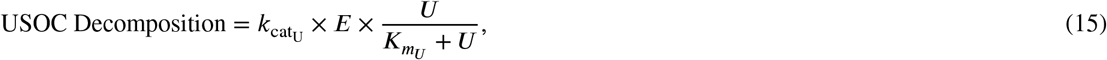

where 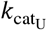, the catalytic constant for USOC decomposition, and 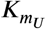, the Michaelis constant for USOC decomposition, are defined (similar to Eqs. (7) and (8)) as

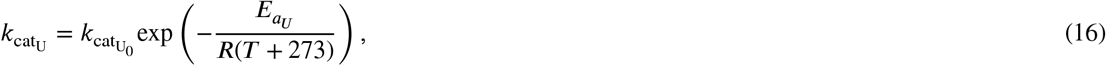

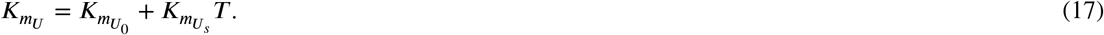

In Eqs. (16)-(17), 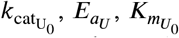, and 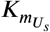respectively represent the pre-exponential coefficient, the activation energy needed to convert substrate into product, the value of 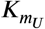 at 0° Celsius, and the slope of the linear function of 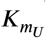.

Substitution of Eqs. (3) and (15) into Eq. (14), and denoting USOC external input as *I*_U_, gives the final form of the ODE for uncontaminated soil organic carbon as

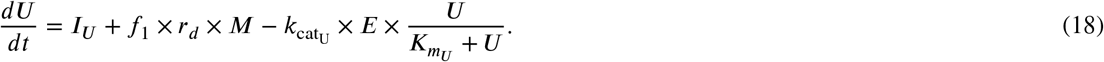

Similar to USOC, contaminated soil organic carbon, CSOC, increases due to external inputs as well as contributions from a fraction of dead microbial biomass, and decreases due to losses from CSOC decomposition. The ODE describing the change of CSOC in time is given as

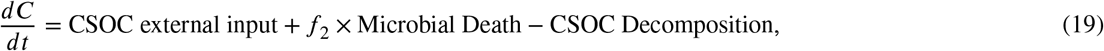

where *C*, as above, denotes the contaminated soil organic carbon, CSOC, and *f*_2_ represents the fraction of dead microbial biomass which is added to the CSOC. The equation for microbial death is given by Eq. (3), and in general, *f*_2_ may differ from *f*_1_.

CSOC decomposition is catalysed via extracellular enzymes and is modeled with Michaelis-Menten kinetics so that

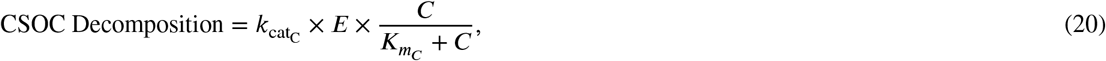

where 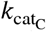, the catalytic constant for CSOC decomposition, and 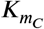, the Michaelis constant for CSOC decomposition, are defined (similar to Eqs. (7) and (8)) as

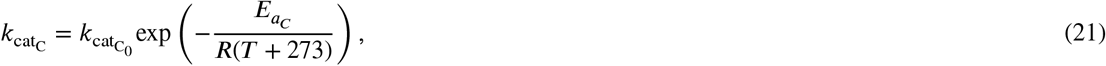

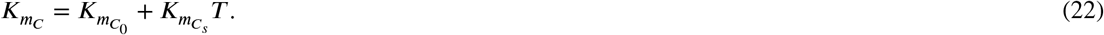

In Eqs. (21)-(22), 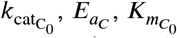, and 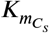 respectively represent the pre-exponential coefficient, the activation energy needed to convert substrate into product, the value of *K* at 0° Celsius, and the slope of the linear function of 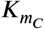.

Substitution of Eqs. (3) and (20) into Eq. (19), and denoting CSOC external input as *I*_*C*_ ODE for contaminated soil organic carbon as, gives the final form of the

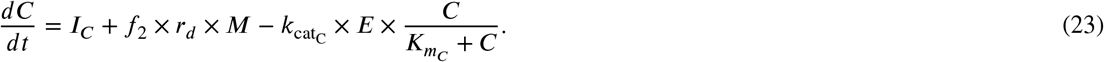

Lastly, dissolved organic carbon, DOC, increases due to external inputs, the remaining 1 −*f*_1_ −*f*_2_ fraction of dead microbial biomass, dead extracellular enzymes, and decomposition of both USOC and CSOC, and decreases due to DOC assimilation. The ODE describing the change of DOC in time is given as

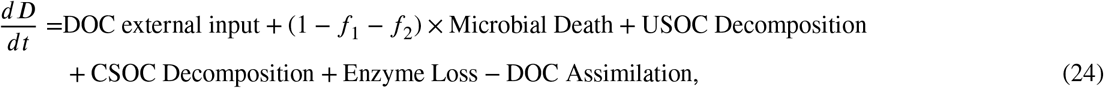

where *D*, as above, denotes dissolved organic carbon, DOC. Substitution of Eqs. (2)-(3), (12), (15) and (20) into Eq. (24), and denoting DOC external input as *I*_*D*_, gives the final form of the ODE for contaminated dissolved organic carbon as

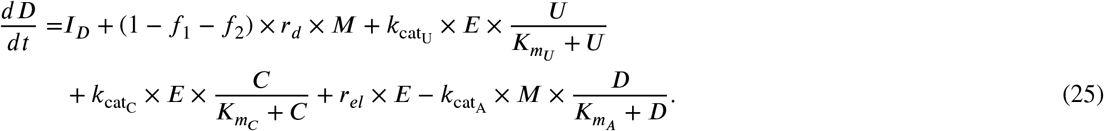

Equations (6), (10), (13), (18), (23), and (25) comprise the contaminated soil model. Given initial conditions and parameter values (Table 1), the ODE system is solved numerically over a 30 year time frame using the ode45 Runge-Kutta solver in MATLAB. Recall that it is possible for CSOC to increase while USOC remains relatively unchanged.

**Table 1.** Initial values and parameter values for the contaminated and uncontaminated soil models in barren and vegetated environments.

| Variable | Units | Value | Citation |
| --- | --- | --- | --- |
| Time, $t$ | Hours, $h$ | 30*8760 | Selected |
| $C(t=0)$ | $mg \cdot cm^{-3}$ | Varies | Selected |
| $U(t=0)$ | $mg \cdot cm^{-3}$ | 100 | Allison et al. (2010) |
| $D(t=0)$ | $mg \cdot cm^{-3}$ | 0.5 | Allison et al. (2010) |
| $M(t=0)$ | $mg \cdot cm^{-3}$ | 0.5 | Allison et al. (2010) |
| $E(t=0)$ | $mg \cdot cm^{-3}$ | 0.01 | Allison et al. (2010) |
| $CO_2(t=0)$ | $mg \cdot cm^{-3}$ | 0 | Selected |
| $r_d$ | $h^{-1}$ | 0.0002 | Allison et al. (2010) |
| $r_{ep0}$ | $h^{-1}$ | 0.000005 | Allison et al. (2010) |
| $r_{ep}$ | $h^{-1}$ | Varies | Calculated (Eq. (5)) |
| $r_{el}$ | $h^{-1}$ | 0.001 | Allison et al. (2010) |
| $f_1$ | Dimensionless | 0.5 | Selected |
| $f_2$ | Dimensionless | 0 | Selected |
| $R$ | $kJ \cdot mol^{-1} \cdot K^{-1}$ | 0.008314 | Mohr, Newell, Taylor and Tiesinga (2025) |
| $T$ | Celsius, $^{\circ}C$ | 20 | Selected |
| $CUE_0$ | Dimensionless | 0.63 | Allison et al. (2010) |
| $CUE_s$ | $^{\circ}C^{-1}$ | -0.016 | Allison et al. (2010) |
| $CUE$ | Dimensionless | 0.31 | Calculated (Eq. (9)) |
| $k_{catA_0}$ | $h^{-1}$ | 100000000 | Allison et al. (2010) |
| $E_{aA}$ | $kJ \cdot mol^{-1}$ | 47 | Allison et al. (2010) |
| $k_{catA}$ | $h^{-1}$ | 0.4176 | Calculated (Eq. (7)) |
| $K_{mA_0}$ | $mg \cdot cm^{-3}$ | 0.1 | Allison et al. (2010) |
| $K_{mA_s}$ | $mg \cdot cm^{-3} \cdot ^{\circ}C^{-1}$ | 0.01 | Allison et al. (2010) |
| $K_{mA}$ | $mg \cdot cm^{-3}$ | 0.3 | Calculated (Eq. (8)) |
| $k_{catU_0}$ | $h^{-1}$ | 100000000 | Allison et al. (2010) |
| $E_{aU}$ | $kJ \cdot mol^{-1}$ | 47 | Allison et al. (2010) |
| $k_{catU}$ | $h^{-1}$ | 0.4176 | Calculated (Eq. (16)) |
| $K_{mU_0}$ | $mg \cdot cm^{-3}$ | 500 | Allison et al. (2010) |
| $K_{mU_s}$ | $mg \cdot cm^{-3} \cdot ^{\circ}C^{-1}$ | 5 | Allison et al. (2010) |
| $K_{mU}$ | $mg \cdot cm^{-3}$ | 600 | Calculated (Eq. (17)) |
| $k_{catC_0}$ | $h^{-1}$ | 1000000 | Selected |
| $E_{aC}$ | $kJ \cdot mol^{-1}$ | 47 | Selected |
| $k_{catC}$ | $h^{-1}$ | 0.004176 | Calculated (Eq. (21)) |
| $K_{mC_0}$ | $mg \cdot cm^{-3}$ | 500 | Selected |
| $K_{mC_s}$ | $mg \cdot cm^{-3} \cdot ^{\circ}C^{-1}$ | 5 | Selected |
| $K_{mC}$ | $mg \cdot cm^{-3}$ | 600 | Calculated (Eq. (22)) |
| $I_C$ | $mg \cdot cm^{-3} \cdot h^{-1}$ | Varies | Selected |
| $I_U$ | $mg \cdot cm^{-3} \cdot h^{-1}$ | Varies | Selected |
| $I_D$ | $mg \cdot cm^{-3} \cdot h^{-1}$ | Varies | Selected |

In this scenario, after some amount of time has elapsed, the CSOC to USOC ratio may satisfy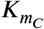. From Eq. (5), this implies that *r*_*ep*_ ≤ 0. If this occurs, we stop the simulation as soon as *r*_*ep*_ crosses the zero ^*U*^threshold. In order to focus on the role of contamination, we have selected the temperature to be *T* = 20°*C* for all simulations.

By adjusting the external inputs, the model can be used to improve our understanding of contamination in both barren and vegetated soils. In particular, in barren environments, the absence of plants means that little/no plant material decomposes on the soil surface, resulting in less organic matter entering the soil. Additionally, plants typically release sugars and other molecules into the soil, but barren soil lacks these inputs as well. Therefore, in our simulations, we estimated the external inputs to USOC and DOC in barren soil to be 100 times lower than in vegetated soil (Luo, Feng, Luo, Baldock and Wang, 2017).

By eliminating the CSOC compartment and the external input to CSOC, Eqs. (6), (10), (13), (18), and (25) comprise the uncontaminated soil model. In the uncontaminated model, the fraction, *f*_2_ of dead microbial biomass added to CSOC is set to zero, and, since *C* = 0 in the uncontaminated model, the enzyme production rate 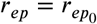(Eq. (5)).

In our uncontaminated model simulations, the dead microbial biomass is equally distributed between DOC and USOC. As with the contaminated model, the uncontaminated model can be used for barren or vegetated environments by adjusting the external inputs to USOC and DOC in barren soil to be 100 times lower than in vegetated soil (Luo et al., 2017).

## 3. Results

We have developed a novel soil contamination model that enables the study of how a soil system containing CSOC, USOC, DOC, microbial biomass, extracellular enzymes, and microbial CO_2_ respiration evolves in time. The primary goal of this work is to study how soil contamination affects basic soil function in vegetated and barren environments over a period of 30 years. In Sec. 3.1, we simulate our microbial-enzyme soil model with and without contamination in both vegetated and barren environments. In Sec. 3.2, we investigate how contamination input rates affect the soil system in both environments. In Sec. 3.3, we examine the impact of press and pulse contamination on soil function in both vegetated and barren environments. Lastly, in Sec. 4, we discuss the implications of the results.

### 3.1. Contamination in vegetated and barren environments

The soil model given by Eqs. (6), (10), (13), (18), (23), and (25), is solved numerically to study the role of press contamination in vegetated and barren soil environments. The initial contamination amount, *C*(*t* = 0) = 0 *mg* ⋅ *cm*^−3^, and the CSOC external input, *I*_*C*_, is given as *I*_*C*_ = 0.0005 *mg* ⋅ *cm*^−3^ ⋅ *h*^−1^ in both the vegetated and barren environments. In the vegetated environment, the USOC and DOC external inputs, *I*_*U*_ and *I*_*D*_, are given as *I*_*U*_ = *I*_*D*_ = 0.0005 *mg* ⋅ *cm*^−3^ ⋅ *h*^−1^, while in the barren environment the USOC and DOC external inputs are given as *I*_*U*_ = *I*_*D*_ = 0.0005/100 *mg* ⋅ *cm*^−3^ ⋅ *h*^−1^ (Luo et al., 2017). We assume that the dead microbial biomass is added equally to the USOC and DOC (*f*_1_ = 0.5), so that no dead microbial biomass is added to the CSOC (*f*_2_ = 0). Because CSOC is significantly more resistant to enzymatic degradation than USOC (Martinho, Abreu, Oliveira, Alegria and Pampulha, 2013; Straube, Nestler, Hansen, Ringleberg, Pritchard and Jones-Meehan, 2003), the catalytic constant for CSOC decomposition, 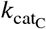, is assumed to be 100 times smaller than the catalytic constant for USOC decomposition so that 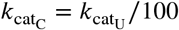.

For comparison, we also considered the uncontaminated soil model found by eliminating Eq. (23) and letting *C* = 0 and *I*_*C*_ = 0. As in the contamination model described above, in the vegetated environment, the USOC and DOC external inputs, *I*_*U*_ and *I*_*D*_, are given as *I*_*U*_ = *I*_*D*_ = 0.0005 *mg* ⋅ *cm*^−3^ ⋅ *h*^−1^, while in the barren environment the USOC and DOC external inputs are given as *I*_*U*_ = *I*_*D*_ = 0.0005/100 *mg* ⋅ *cm*^−3^ ⋅ *h*^−1^. We also maintain *f*_1_ = 0.5 and *f*_2_ = 0 as was used in the contaminated model. Since *C* = 0, the enzyme production rate 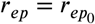 (Eq. (5)).

We numerically solve the system of model equations over 30 years and visualize the time evolution of CSOC, USOC, DOC, microbial biomass, extracellular enzymes, and CO_2_ for the uncontaminated and contaminated microbialenzyme soil system in vegetated and barren environments (Figures 2 and 3). In the contaminated, barren system, the amount of CSOC becomes larger than the amount of USOC after approximately 22.6 years. Once CSOC becomes larger than USOC, the enzyme production rate *r*_*ep*_ ≤ 0 and we stop the simulation at that moment in time (Figures 2 (a), (c) and (e) and 3 (b), (d) and (f)). The amounts of each quantity for each scenario at the end of the simulated time period are shown in Table 2.

**Table 2.** Final amounts of CSOC, *C* (*mg* ⋅ *cm*^−3^), USOC, *U* (*mg* ⋅ *cm*^−3^), DOC, *D* (*mg* ⋅ *cm*^−3^), microbial biomass, *M* (*mg* ⋅ *cm*^−3^), extracellular enzymes, *E* (*mg* ⋅ *cm*^−3^), and CO_2_ (*mg* ⋅ *cm*^−3^) for uncontaminated and contaminated microbial-enzyme models in vegetated and barren environments. In the contaminated, barren system, the amount of CSOC becomes larger than the amount of USOC after 22.6119 years, the amount of extracellular enzymes is effectively zero (3.1101e-09 *mg* ⋅ *cm*^−3^), and the simulation is stopped. For the other three scenarios, the simulation is stopped after 30 years.

|  | *C* |  | *U* |  | *D* |  |
| --- | --- | --- | --- | --- | --- | --- |
|  | Vegetated | Barren | Vegetated | Barren | Vegetated | Barren |
| Uncontaminated | — | — | 111.533 | 97.798 | 0.0004760 | 0.0004759 |
| Contaminated | 131.054 | 99.035 | 188.350 | 98.546 | 0.0004662 | 0.0004587 |
|  | *M* |  | *E* |  | CO<sub>2</sub> |  |
|  | Vegetated | Barren | Vegetated | Barren | Vegetated | Barren |
| Uncontaminated | 2.182 | 0.019 | 0.011 | 0.0000948 | 250.084 | 5.820 |
| Contaminated | 1.273 | 0.009 | 0.002 | 3.1101e-09 | 174.531 | 4.444 |

**Figure 2.**
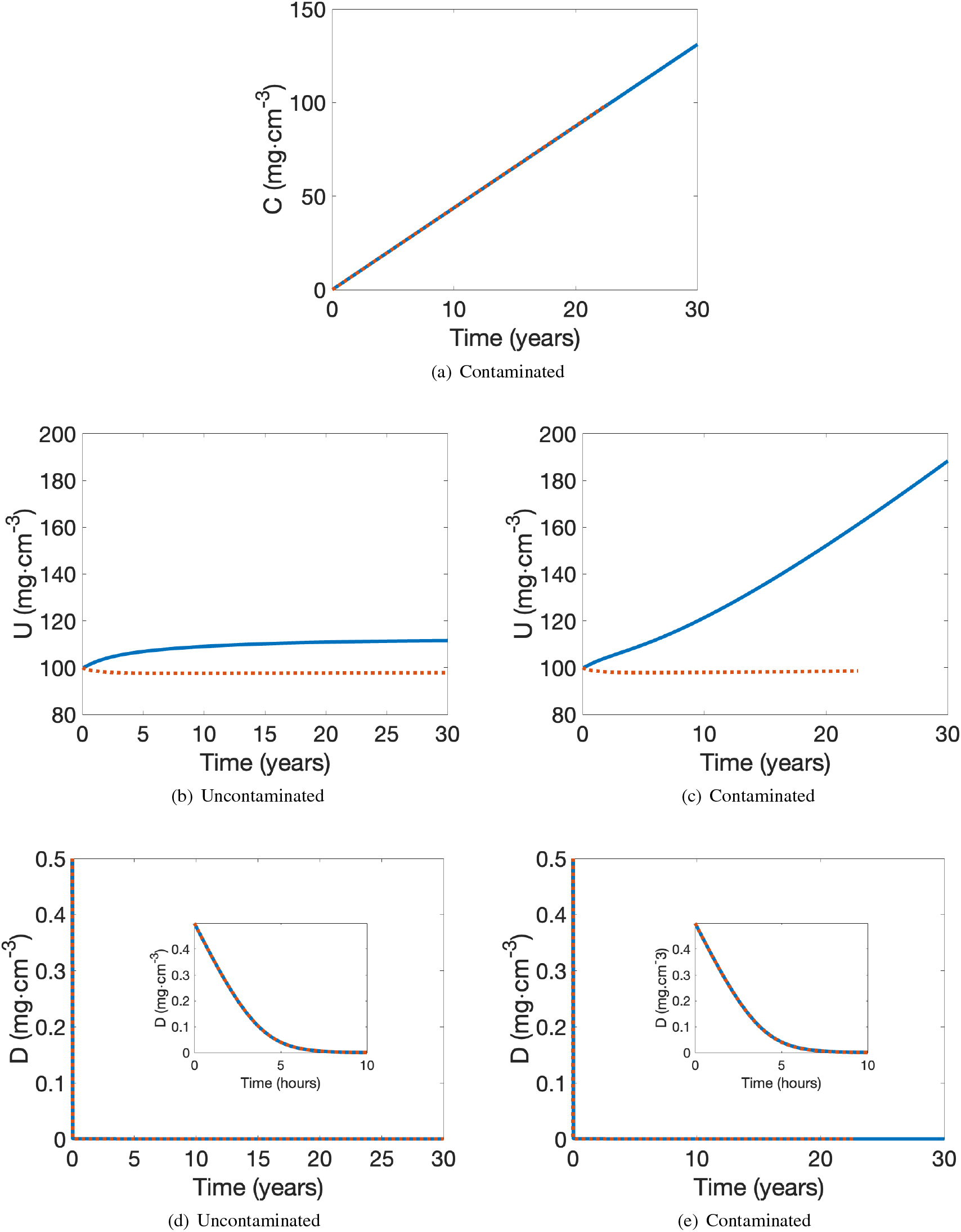
Comparison of (a) CSOC, *C* (*mg* ⋅ *cm*^−3^), (b,c) USOC, *U* (*mg* ⋅ *cm*^−3^), and (d,e) DOC, *D* (*mg* ⋅ *cm*^−3^) for uncontaminated and contaminated microbial-enzyme models in vegetated (continuous, blue curve) and barren (dashed, orange curve) environments. The figures are arranged in columns to facilitate the comparison of uncontaminated (b,d) and contaminated (a,c,e) conditions. We assume press contamination with *I*_*C*_ = 0.0005 *mg* ⋅ *cm*^−3^ ⋅ *h*^−1^. In the contaminated, barren system, the amount of CSOC becomes larger than the amount of USOC after approximately 22.6 years, and the simulation is stopped.

**Figure 3.**
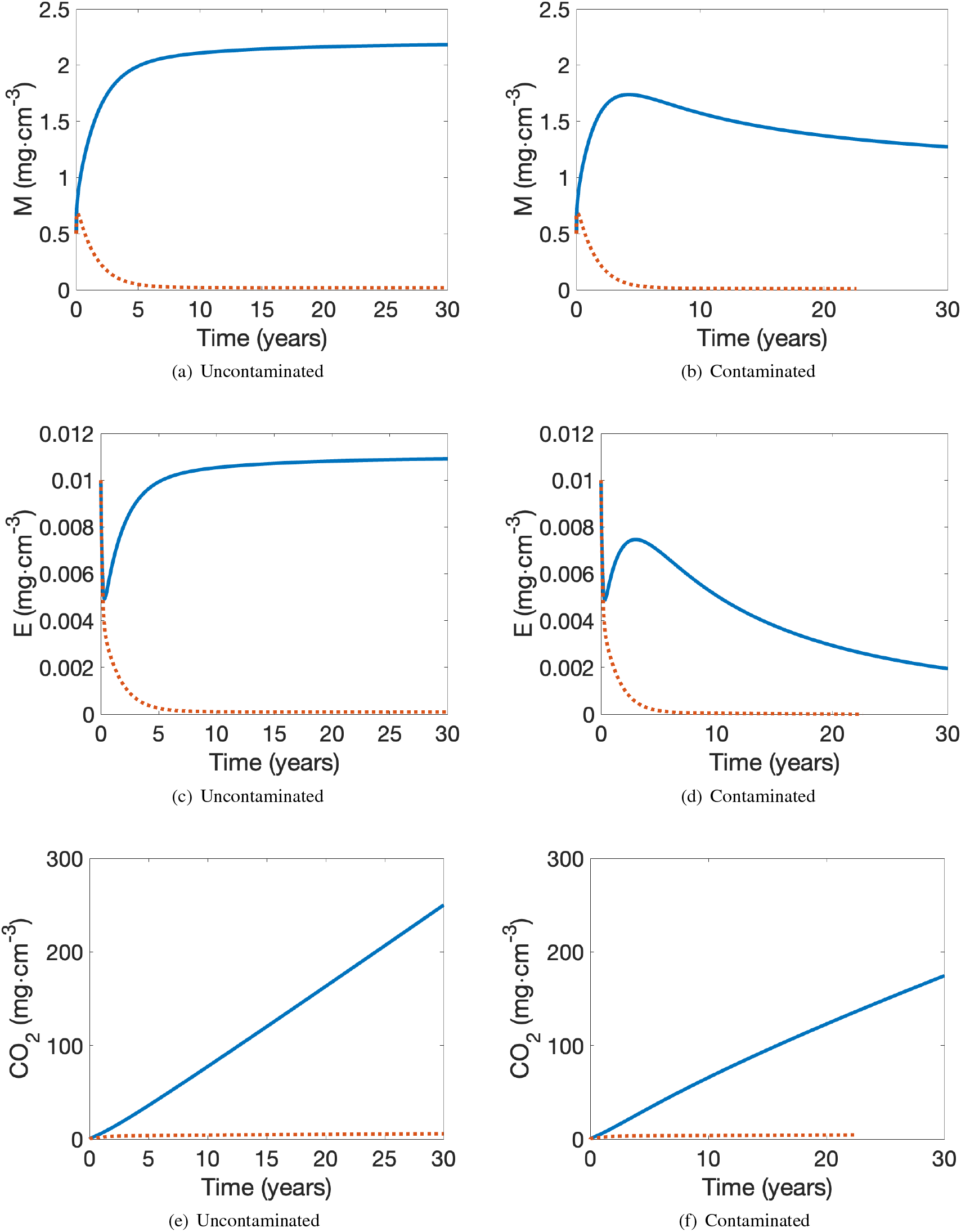
Comparison of (a,b) microbial biomass, *M* (*mg* ⋅ *cm*^−3^), (c,d) extracellular enzymes, *E* (*mg* ⋅ *cm*^−3^), and (e,f) CO_2_ (*mg* ⋅ *cm*^−3^) for uncontaminated and contaminated microbial-enzyme models in vegetated (continuous, blue curve) and barren (dashed, orange curve) environments. The figures are arranged in columns to facilitate the comparison of uncontaminated (a,c,e) and contaminated (b,d,f) conditions. *I*_*C*_ = 0.0005 *mg* ⋅ *cm*^−3^ ⋅ *h*^−1^. In the contaminated, barren system, the amount of CSOC becomes larger than the amount of USOC after approximately 22.6 years, and the simulation is stopped.

The time evolution of CSOC shows a similar trend in both vegetated and barren environments (Figure 2 (a)). Because *f*_2_ = 0 and 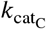 (Table 1) and *E* (Figure 3 (d)) are relatively small, one can see from Eq. (23) that CSOC will increase approximately linearly in time with slope given by *I*_*C*_. There is a small difference in the amount of extracellular enzymes, *E*, between the vegetated and barren environments. This difference will lead to a very small change in the time evolution of CSOC. However, because of the long time scale which has been used in Figure 2 (a), the two curves appear to overlap one another.

The time evolution of USOC in the barren environment is similar for both contaminated and uncontaminated systems (Figures 2 (b)-(c)). In both scenarios, there is initially a small decrease in USOC because the amount of USOC decomposition is larger than the combined addition of USOC from dead microbial biomass and the external input (Eq. 18). As the amount of microbial biomass (Figures 3 (a)-(b)) and extracellular enzyme (Figures 3 (c)-(d)) decrease to near zero, then one can see from Eq. (18) that USOC will increase approximately linearly in time with slope given by *I*_*U*_. Because *I*_*U*_ in the barren environment is so small, USOC will only increase about 1.31 *mg* ⋅ *cm*^−3^ over a 30 year span. This leads to USOC time evolution curves which appear flat (Figures 2 (b)-(c)). In the vegetated environment, contamination leads to an overall decline in microbial biomass and extracellular enzymes when compared to the uncontaminated scenario. The larger amount of enzyme activity in the vegetated environment is consistent with the increases in enzyme activity due to revegetation that have been reported during restoration of desertified soils (Zhang, Chen, Chen, Tan, Duan, Wu, Li and Fan, 2015). In the vegetated environment, continued external carbon inputs maintain greater amounts of USOC throughout the simulation. This prediction is consistent with field studies showing that vegetation of sandy barren lands increased SOC pools through sustained vegetation inputs (Moharana, Naitam, Verma, Meena, Kumar, Tailor, Singh, Singh and Samal, 2017). The enzyme decline due to contamination leads to an order of magnitude decrease in USOC decomposition over the 30 year time frame. The end result is that USOC increases much more in contaminated conditions than in uncontaminated soil (Figures 2 (b)-(c)).

The DOC for both uncontaminated and contaminated conditions show similar trends over the 30 years in vegetated and barren environments (Figures 2 (d)-(e)). Also, the DOC values drop sharply to near zero within the first five hours (insets of Figures 2 (d)-(e)). In a barren environment, there is a decay over time of both microbial biomass and enzymes.

Because of this, all terms in Eq. (25) remain very small, and there is no way for the DOC to recover. In a vegetated environment, one can see that the number of microbes increases rapidly at the beginning, indicating that they are consuming a large amount of DOC to grow (Figures 3 (a)-(b)). At the start, DOC assimilation is much larger than the sum of all the other terms affecting the growth of DOC which leads to an initial drop in DOC concentration. As time evolves, all terms in Eq. (25) are small and the overall rate of change of DOC is near zero. As in the barren environment, DOC cannot recover (Figures 2 (d)-(e)). Importantly, because the DOC external input differs in the vegetated and barren environments, there are small differences in the DOC amounts. These differences cannot be seen at the scale shown in Figures 2 (d)-(e) but do affect the microbial growth in these two environments.

Microbial activity initially increases in both the barren and vegetated environments for both uncontaminated and contaminated scenarios (Figures 3 (a)-(b)). This is because at the start, DOC assimilation is larger than the losses due to microbial death and enzyme production (Eq. (6)). After a short period of time, the previously mentioned differences in DOC amounts in the barren and vegetated environments leads to a difference in the microbial activity. While the amount of microbes continues to increase in the vegetated environment, in the barren environment the microbial activity begins to decrease, and ultimately collapses to near zero in both uncontaminated and contaminated scenarios. In the vegetated environment, microbial growth slows down and ultimately grows very slowly in uncontaminated conditions. However, in contaminated conditions, the contamination affects enzyme production (Eqs. (4)-(5)) which ultimately leads to a turnover and slow decay of microbial activity in the vegetated environment.

Enzyme activity initially decreases in both the barren and vegetated environments for both uncontaminated and contaminated scenarios (Figures 3 (c)-(d)). This is because at the start, enzyme loss is larger than enzyme production (Eq. (13)). In the barren environment for both uncontaminated and contaminated scenarios, microbial activity starts to decline after a short period of time. This causes a continued decline in enzyme activity, which ultimately collapses to near zero. In the vegetated, uncontaminated environment, after some time, microbial growth increases enough so that enzyme production is greater than enzyme loss and the amount oof enzymes starts to grow. This rapid enzyme growth slows down and continues to grow slowly as the microbial growth slows down. Similarly, in the vegetated, contaminated environment, enzyme activity increases as microbial activity increases; ultimately, enzyme activity will decline, following the decline of the microbial population.

Figures 3 (e)-(f) show the cumulative amount of CO_2_ production. Because the amount of DOC is near zero for all environments, CO_2_ production is an approximately linear function of *M* (Eq. 10). Because microbial activity is very low in barren environments, little CO_2_ accumulates. In contrast, CO_2_ does accumulate in vegetated environments, with a larger accumulation in uncontaminated soils since there is a larger microbial population in this scenario.

### 3.2. The role of contamination input rates

To understand how different amounts of contamination affects the time evolution of CSOC, USOC, DOC, microbial biomass, extracellular enzymes, and CO_2_, the soil model described by Eqs. (6), (10), (13), (18), (23), and (25) is numerically solved for varying levels of external CSOC contamination input rates. The initial contamination level is set to *C*(*t* = 0) = 0 *mg* ⋅ *cm*^−3^, and the model is simulated for five choices of external CSOC contamination input rates,*I*_*C*_ = {0.0001, 0.0002, 0.0003, 0.0004, 0.0005} *mg* ⋅ *cm*^−3^ ⋅ *h*^−1^ in both the vegetated and barren environments.

As in Section 3.1, in the vegetated environment, the USOC and DOC external inputs, *I*_*U*_ and *I*_*D*_, are given as *I*_*U*_ = *I*_*D*_ = 0.0005 *mg* ⋅ *cm*^−3^ ⋅ *h*^−1^, while in the barren environment the USOC and DOC external inputs are given as *I*_*U*_ = *I*_*D*_ = 0.0005/100 *mg* ⋅ *cm*^−3^ ⋅ *h*^−1^. Dead microbial biomass is added equally to the USOC and DOC (*f*_1_ = 0.5), so that no dead microbial biomass is added to the CSOC (*f*_2_ = 0), and the catalytic constant for CSOC decomposition, 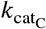, is assumed to be 100 times smaller than the catalytic constant for USOC decomposition so that 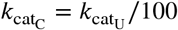. We numerically solve the system of model equations over 30 years and visualize the time evolution of CSOC, USOC, DOC, microbial biomass, extracellular enzymes, and CO_2_ for the contaminated microbial-enzyme soil system in vegetated and barren environments for five choices of external CSOC contamination input rates (Figures 4 and 5). In the contaminated, barren system, the amount of CSOC becomes larger than the amount of USOC after approximately 22.6 years when *I*_*c*_ = 0.0005 *mg* ⋅ *cm*^−3^ ⋅ *h*^−1^ and after approximately 28.3 years when *I*_*c*_ = 0.0004 *mg* ⋅ *cm*^−3^ ⋅ *h*^−1^. Once CSOC becomes larger than USOC, the enzyme production rate *r*_*ep*_ ≤ 0 and we stop the simulation at that moment in time (Figures 4 (b), (d) and (f) and 5 (b), (d) and (f)).

**Figure 4.**
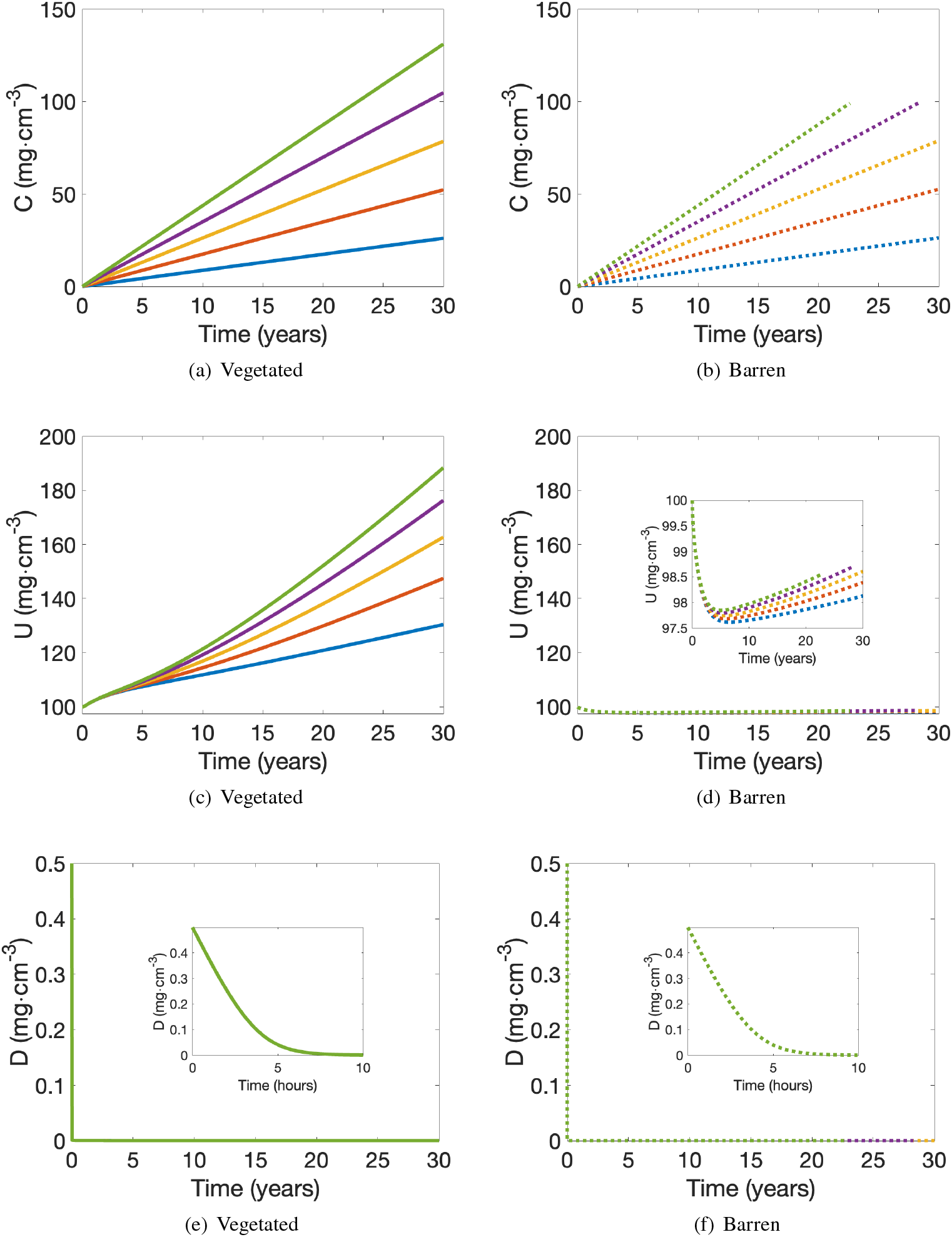
Comparison of (a,b) CSOC, *C* (*mg* ⋅ *cm*^−3^), (c,d) USOC, *U* (*mg* ⋅ *cm*^−3^), and (e,f) DOC (*mg* ⋅ *cm*^−3^) for the contaminated microbial-enzyme model in vegetated (a,c,e) and barren (b,d,f) environments. Five choices of external CSOC contamination input rates, *I*_*C*_ = 0.0001 (blue curve), *I*_*C*_ = 0.0002 (red curve), *I*_*C*_ = 0.0003 (yellow curve), *I*_*C*_ = 0.0004 (purple curve), and *I*_*C*_ = 0.0005 (green curve) *mg* ⋅ *cm*^−3^ ⋅ *h*^−1^ are considered for vegetated (continuous curves) and barren (dashed curves) environments. The figures are arranged in columns to facilitate the comparison of vegetated (a,c,e) and barren (b,d,f) environments. In the barren system, the amount of CSOC becomes larger than the amount of USOC after approximately 22.6 years when *I*_*c*_ = 0.0005 *mg*⋅*cm*^−3^ ⋅*h*^−1^ and after approximately 28.3 years when *I*_*c*_ = 0.0004 *mg*⋅*cm*^−3^ ⋅*h*^−1^, and the simulation is stopped.

**Figure 5.**
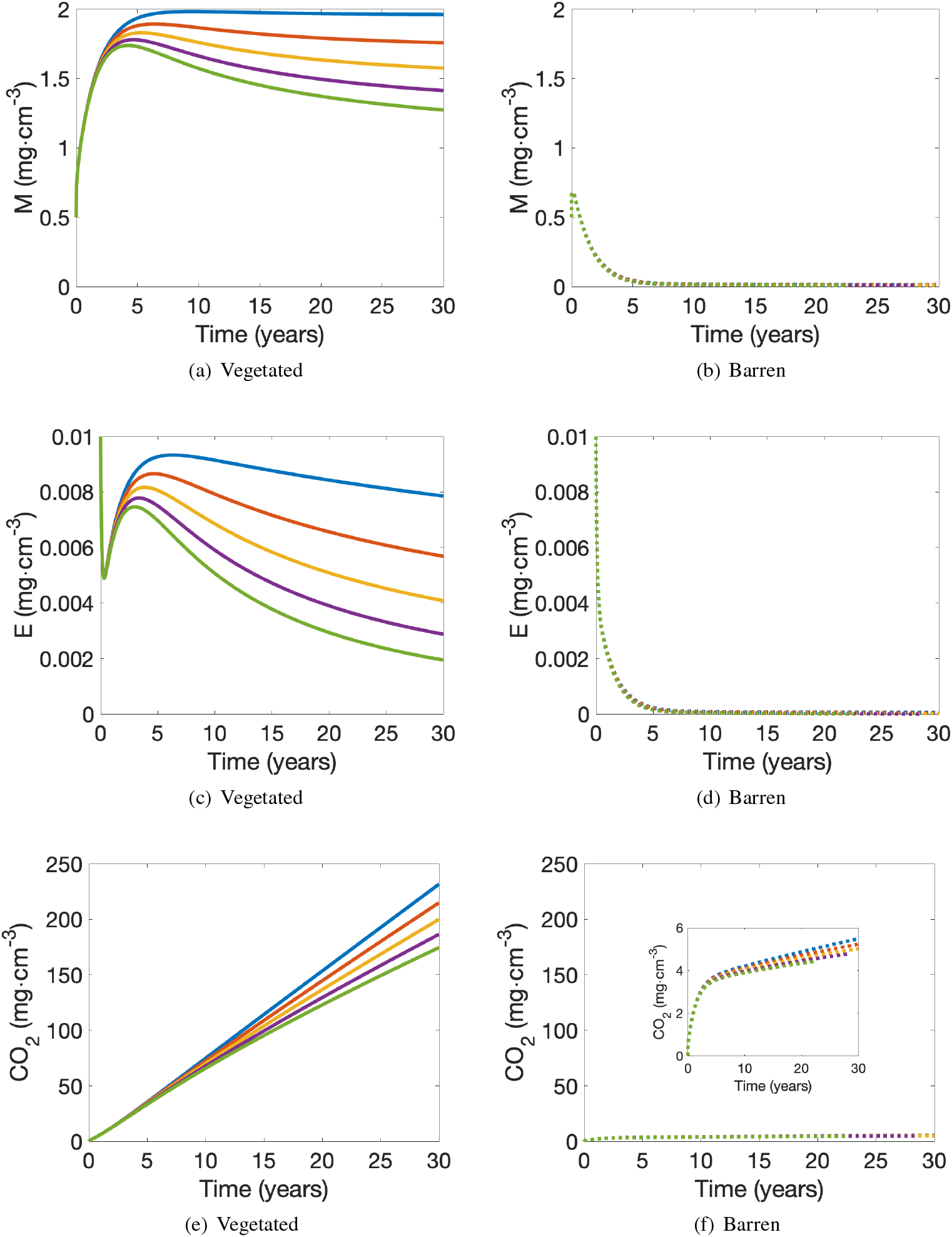
Comparison of (a,b) microbial biomass, *M* (*mg* ⋅ *cm*^−3^), (c,d) extracellular enzymes, *E* (*mg* ⋅ *cm*^−3^), and (e,f) CO_2_ (*mg* ⋅ *cm*^−3^) for the contaminated microbial-enzyme model in vegetated (a,c,e) and barren (b,d,f) environments. Five choices of external CSOC contamination input rates, *I*_*C*_ = 0.0001 (blue curve), *I*_*C*_ = 0.0002 (red curve), *I*_*C*_ = 0.0003 (yellow curve), *I*_*C*_ = 0.0004 (purple curve), and *I*_*C*_ = 0.0005 (green curve) *mg* ⋅ *cm*^−3^ ⋅ *h*^−1^ are considered for vegetated (continuous curves) and barren (dashed curves) environments. The figures are arranged in columns to facilitate the comparison of vegetated (a,c,e) and barren (b,d,f) environments. In the barren system, the amount of CSOC becomes larger than the amount of USOC after approximately 22.6 years when *I*_*c*_ = 0.0005 *mg* ⋅ *cm*^−3^ ⋅ *h*^−1^ and after approximately 28.3 years when *I*_*c*_ = 0.0004 *mg* ⋅ *cm*^−3^ ⋅ *h*^−1^, and the simulation is stopped.

The results in Figures 4 and 5 for *I*_*c*_ = 0.0005 *mg* ⋅ *cm*^−3^ ⋅ *h*^−1^ (continuous, green curves for the vegetated environment and dashed, green curves for the barren environment) are identical to the results for the contaminated soil-system shown in Figures 2 (a,c,e) and 3 (b,d,f) (continuous, blue curves for the vegetated environment and dashed,orange curves for the barren environment). All other curves in Figures 4 and 5 show how a change in the amount of external press contamination affects the time evolution of CSOC, USOC, DOC, microbial biomass, extracellular enzymes, and CO_2_.

The qualitative dynamics of each quantity is generally the same for the different contamination rates (e.g. CSOC exhibits approximately linear growth, DOC exhibits rapid decay, etc.). There are, however, quantitative differences in the growth and decay rates and the values at the end of the simulation time period (Table 3). The explanation of the dynamical behaviour is the same as was described for each quantity in Section 3.1.

**Table 3.** Final amounts of CSOC, *C* (*mg* ⋅ *cm*^−3^), USOC, *U* (*mg* ⋅ *cm*^−3^), DOC, *D* (*mg* ⋅ *cm*^−3^), microbial biomass, *M* (*mg* ⋅ *cm*^−3^), extracellular enzymes, *E* (*mg* ⋅ *cm*^−3^), and CO_2_ (*mg* ⋅ *cm*^−3^) for the contaminated microbial-enzyme model in vegetated and barren environments for five choices of external CSOC contamination input rates, *I*_*C*_ = {0.0001, 0.0002, 0.0003, 0.0004, 0.0005} *mg* ⋅ *cm*^−3^ ⋅ *h*^−1^. In the contaminated, barren system, with *I*_*C*_ = 0.0004, the amount of CSOC becomes larger than the amount of USOC after 28.282 years, the amount of extracellular enzymes is effectively zero (1.0629e-09 *mg* ⋅*cm*^−3^), and the simulation is stopped. Similarly, in the contaminated, barren system, with *I*_*C*_ = 0.0005, the amount of CSOC becomes larger than the amount of USOC after 22.6119 years, the amount of extracellular enzymes is effectively zero (3.0588e-09 *mg* ⋅ *cm*^−3^), and the simulation is stopped. For all other scenarios, the simulation is stopped after 30 years.

|  | *C* |  | *U* |  | *D* |  |
| --- | --- | --- | --- | --- | --- | --- |
| *I<sub>C</sub>* | Vegetated | Barren | Vegetated | Barren | Vegetated | Barren |
| 0.0001 | 26.084 | 26.278 | 130.408 | 98.132 | 0.0004738 | 0.0004708 |
| 0.0002 | 52.258 | 52.557 | 147.459 | 98.395 | 0.0004714 | 0.0004665 |
| 0.0003 | 78.490 | 78.837 | 162.714 | 98.611 | 0.0004698 | 0.0004625 |
| 0.0004 | 104.759 | 99.096 | 176.291 | 98.703 | 0.0004678 | 0.0004593 |
| 0.0005 | 131.054 | 99.035 | 188.350 | 98.546 | 0.0004662 | 0.0004587 |
|  | *M* |  | *E* |  | CO<sub>2</sub> |  |
| *I<sub>C</sub>* | Vegetated | Barren | Vegetated | Barren | Vegetated | Barren |
| 0.0001 | 1.962 | 0.015 | 0.008 | 0.0000548 | 231.628 | 5.493 |
| 0.0002 | 1.758 | 0.012 | 0.006 | 0.0000288 | 214.890 | 5.234 |
| 0.0003 | 1.574 | 0.010 | 0.004 | 0.0000106 | 199.868 | 5.020 |
| 0.0004 | 1.413 | 0.009 | 0.003 | 1.0629e-09 | 186.464 | 4.778 |
| 0.0005 | 1.273 | 0.009 | 0.002 | 3.0588e-09 | 174.531 | 4.439 |

As expected, with increasing contamination, *I*_*C*_, CSOC increases approximately linearly in time with slope given by *I*_*C*_. Because there is a small difference in the amount of extracellular enzymes, *E*, in the vegetated and barren environments, there will be a very small change in the time evolution of CSOC for a specified value of *I*_*C*_. The entire time evolution and the final amounts of CSOC in the vegetated and barren environments are very close to one another. In the barren environment, the different contamination rates have little effect on USOC, DOC, microbial biomass, extracellular enzymes, and CO_2_. All curves follow the same trajectory for a few years, and then begin to separate. However, the differences at the end of the simulation period are very small (Table 3). In the vegetated environment, as in the barren environment, the curves for USOC, DOC, microbial biomass, extracellular enzymes, and CO_2_ follow the same trajectory for a few years before separating. In contrast to the barren environment, the trajectories and final amounts are significantly different. With increasing contamination rate, *I*_*C*_, one sees an expected decline in DOC,microbial biomass, extracellular enzymes, and CO_2_.

### 3.3. The impact of press and pulse contamination

To understand how press and pulse contamination affects the time evolution of CSOC, USOC, DOC, microbial biomass, extracellular enzymes, and CO_2_, the soil model described by Eqs. (6), (10), (13), (18), (23), and (25) is numerically solved for two types of external CSOC contamination input: (1) continuous, press contamination, and (2) a one-time, discrete pulse contamination event. For press contamination, as used in Sections 3.1 and 3.2, the initial contamination level is set to *C*(*t* = 0) = 0 *mg* ⋅ *cm*^−3^, and the model is simulated with an external CSOC contamination input rates, *I*_*C*_ = 0.0001 *mg* ⋅ *cm*^−3^ ⋅ *h*^−1^ in both the vegetated and barren environments. For the pulse contamination event, the total amount of contamination entering the system over 30 years as press contamination is deposited at the start of the simulation so that *C*(*t* = 0) = 26.28 *mg* ⋅ *cm*^−3^ in both the vegetated and barren environments, with no additional contamination after the initial contaminant deposition.

As in Section 3.1 and 3.2, in the vegetated environment, the USOC and DOC external inputs, *I*_*U*_ and *I*_*D*_, are given as *I*_*U*_ = *I*_*D*_ = 0.0005 *mg* ⋅ *cm*^−3^ ⋅ *h*^−1^, while in the barren environment the USOC and DOC external inputs are given as *I*_*U*_ = *I*_*D*_ = 0.0005/100 *mg* ⋅ *cm*^−3^ ⋅ *h*^−1^. Dead microbial biomass is added equally to the USOC and DOC (*f*_1_ = 0.5), so that no dead microbial biomass is added to the CSOC (*f*_2_ = 0), and the catalytic constant for CSOC decomposition, 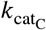, is assumed to be 100 times smaller than the catalytic constant for USOC decomposition so that 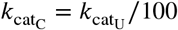.

We numerically solve the system of model equations over 30 years and visualize the time evolution of CSOC, USOC, DOC, microbial biomass, extracellular enzymes, and CO_2_ for the contaminated microbial-enzyme soil system in vegetated and barren environments under pulse and press contamination (Figures 6 and 7). The results in Figures 6 (b,d,f) and 7 (b,d,f) for press contamination with *I*_*c*_ = 0.0001 *mg* ⋅ *cm*^−3^ ⋅ *h*^−1^ are identical to the results for the contaminated soil-system shown in Figures 4 and 5 (continuous, blue curves for the vegetated environment and dashed, blue curves for the barren environment). The results in Figures 6 (a,c,e) and 7 (a,c,e) show how pulse contamination affects the time evolution of CSOC, USOC, DOC, microbial biomass, extracellular enzymes, and CO_2_.

**Figure 6.**
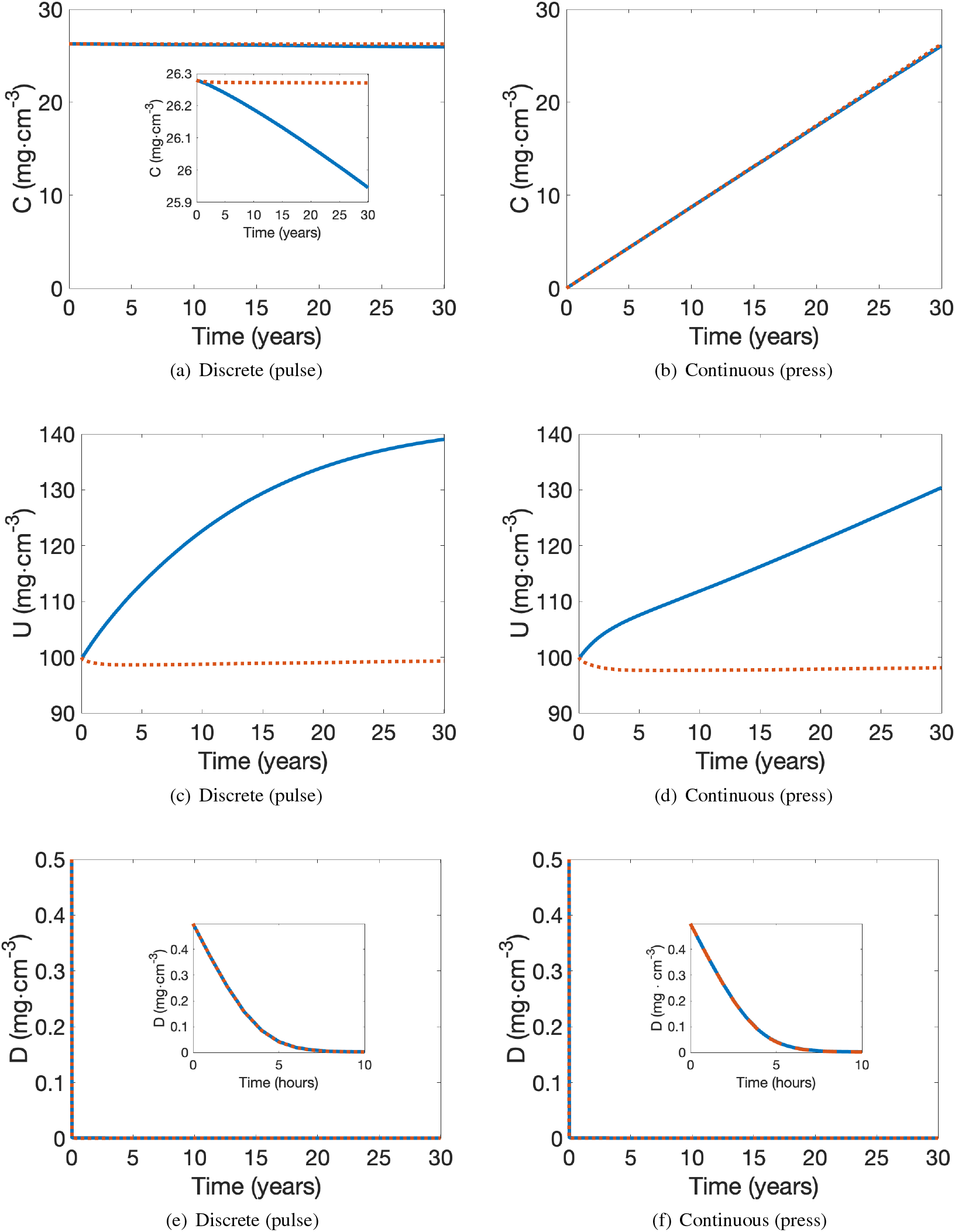
Comparison of (a,b) CSOC, *C* (*mg* ⋅ *cm*^−3^), (c,d) USOC, *U* (*mg* ⋅ *cm*^−3^), and (e,f) DOC, *D* (*mg* ⋅ *cm*^−3^) for the contaminated microbial-enzyme models under discrete (pulse) contamination (a,c,e) and continuous (press) contamination (b,d,f) in vegetated (continuous, blue curve) and barren (dashed, orange curve) environments. The figures are arranged in columns to facilitate the comparison of pulse (a,c,e) and press (b,d,f) contamination. Under press contamination, the contamination input rate is *I*_*C*_ = 0.0001 *mg* ⋅ *cm*^−3^ ⋅ *h*^−1^. Under pulse contamination, the total amount of contamination entering the system over 30 years as press contamination is deposited at the start of the simulation so that *C*(*t* = 0) = 26.28 *mg* ⋅ *cm*^−3^, with no additional contamination after the initial contaminant deposition.

**Figure 7.**
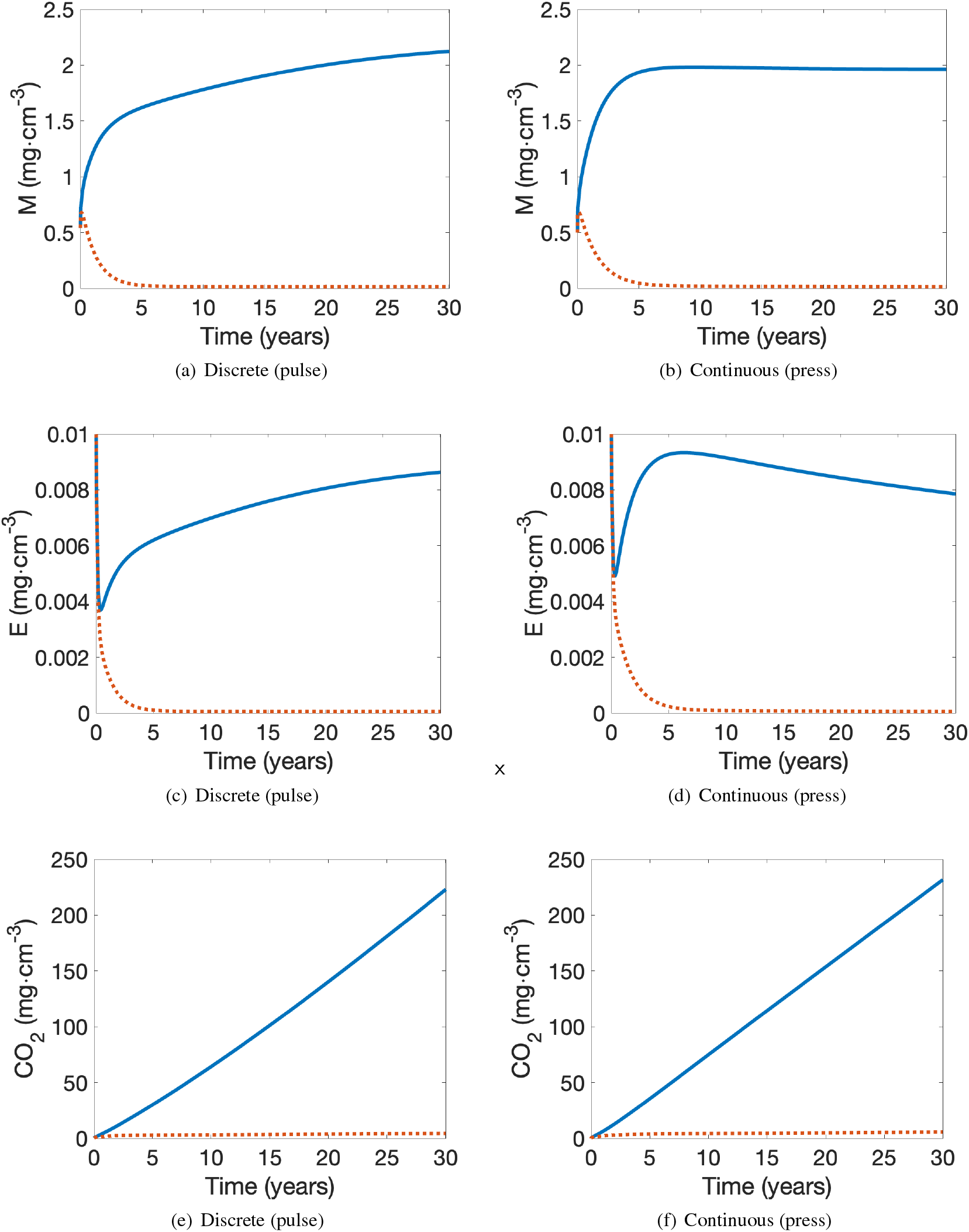
Comparison of (a,b) mirobial biomass, *M* (*mg* ⋅ *cm*^−3^), (c,d) extracellular enzymes, *E* (*mg* ⋅ *cm*^−3^), and (e,f) CO_2_ (*mg* ⋅ *cm*^−3^) for the contaminated microbial-enzyme models under discrete (pulse) contamination (a,c,e) and continuous (press) contamination (b,d,f) in vegetated (continuous, blue curve) and barren (dashed, orange curve) environments. The figures are arranged in columns to facilitate the comparison of pulse (a,c,e) and press (b,d,f) contamination. Under press contamination, the contamination input rate is *I*_*C*_ = 0.0001 *mg* ⋅ *cm*^−3^ ⋅ *h*^−1^. Under pulse contamination, the total amount of contamination entering the system over 30 years as press contamination is deposited at the start of the simulation so that *C*(*t* = 0) = 26.28 *mg* ⋅ *cm*^−3^, with no additional contamination after the initial contaminant deposition.

The qualitative dynamics of each quantity is generally the same for the different types of contamination. There are, however, quantitative differences in the growth and decay rates and the values at the end of the simulation time period (Table 4). The explanation of the dynamical behaviour is the same as was described for each quantity in Section 3.1.

**Table 4.** Final amounts of CSOC, *C* (*mg* ⋅ *cm*^−3^), USOC, *U* (*mg* ⋅ *cm*^−3^), DOC, *D* (*mg* ⋅ *cm*^−3^), microbial biomass, *M* (*mg* ⋅ *cm*^−3^), extracellular enzymes, *E* (*mg* ⋅ *cm*^−3^), and CO_2_ (*mg* ⋅ *cm*^−3^) for the contaminated microbial-enzyme model in vegetated and barren environments under discrete (pulse) and continuous (press) contamination. For all scenarios, the simulation is stopped after 30 years.

|  | *C* |  | *U* |  | *D* |  |
| --- | --- | --- | --- | --- | --- | --- |
|  | Vegetated | Barren | Vegetated | Barren | Vegetated | Barren |
| Pulse | 25.946 | 26.271 | 139.040 | 99.329 | 0.0004749 | 0.0004729 |
| Press | 26.084 | 26.278 | 130.408 | 98.132 | 0.0004738 | 0.0004708 |
|  | *M* |  | *E* |  | CO<sub>2</sub> |  |
|  | Vegetated | Barren | Vegetated | Barren | Vegetated | Barren |
| Pulse | 2.123 | 0.015 | 0.009 | 5.49e-05 | 222.972 | 4.302 |
| Press | 1.962 | 0.015 | 0.008 | 5.48e-05 | 231.628 | 5.493 |

Under pulse contamination, there is a one-time deposition of 26.28 *mg* ⋅ *cm*^−3^ of CSOC, with no additional contamination occurring. After 30 years, in a barren environment, very little of the CSOC has degraded with 26.271 *mg* ⋅ *cm*^−3^ remaining. As expected, in a vegetated environment, there is more degradation of the CSOC, with 25.946 *mg* ⋅ *cm*^−3^ remaining after 30 years. In contrast, under press contamination, the initial amount of CSOC is 0 *mg* ⋅ *cm*^−3^, and over time the amount grows approximately linearly as explained in Section 3.1. After 30 years of growth, the amount of CSOC in the soil is higher than what remains under pulse contamination for both barren and vegetated environments, although the relative differences are less than one percent.

Similarly, the 30 year outcomes are better for all other quantities under pulse contamination. In the vegetated environment, the final amounts of USOC, DOC, microbial biomass, extracellular enzymes are all higher for a one-time pulse contamination event as compared with continuous press contamination (with relative differences again being small). The same is true in the barren environment, except there is no difference in microbial biomass, and because there is little dynamic activity in the barren environment, the relative differences are very small. For both environments, there is less CO_2_ produced under pulse contamination.

## 4. Discussion

The contaminated microbial-enzyme soil model that we have developed enables the study of the temporal dynamics of CSOC, USOC, DOC, microbial biomass, extracellular enzymes, and CO_2_ for different types and amounts of contamination, for different sets of parameter values, and for different lengths of time. In the barren environment, there is little dynamical activity. Even in uncontaminated soil, DOC, microbial biomass, and extracellular enzymes collapse to near zero, with little CO_2_ being respired and USOC remaining nearly constant. Furthermore, over a long time period of 30 years, different types (press and pulse) and amounts of contamination do not significantly affect the amounts of USOC, DOC, microbial biomass, extracellular enzymes or CO_2_ in the barren environment. In the absence of vegetation, contamination has little effect on the soil dynamics.

In contrast, the amount and type of contamination has a significant effect on vegetated soil. Comparison of the amounts of USOC, DOC, microbial biomass, extracellular enzymes and CO_2_ after 30 years for the uncontaminated soil versus the lowest amount of press contamination we considered (*I*_*C*_ = 0.0001 *mg* ⋅ *cm*^−3^ ⋅ *h*^−1^) shows a 16.92% increase in USOC, a 0.46% decrease in DOC, a 10.08% decrease in microbial biomass, a 27.27% decrease in extracellular enzymes, and a 7.38% decrease in CO_2_ when the soil becomes contaminated. A similar comparison of the different amounts after 30 years shows an additional 44.43% increase in USOC, 1.60% decrease in DOC, 35.12% decrease in microbial biomass, 75.00% decrease in extracellular enzymes, and 24.65% decrease in CO_2_ when the soil contamination increases by a factor of 5 from a press contamination rate of *I*_*C*_ = 0.0001 *mg* ⋅ *cm*^−3^ ⋅ *h*^−1^ to *I*_*C*_ = 0.0005 *mg* ⋅ *cm*^−3^ ⋅ *h*^−1^. This underscores the role of vegetation in promoting soil carbon cycling, leading to increased microbial activation, higher CO_2_ release, greater USOC accumulation, and more enzyme production under varying contamination concentration in the soil system.

Importantly, the changes in USOC, DOC, microbial biomass, extracellular enzymes and CO_2_ with changing contamination rates is not linear. The greatest changes in the five quantities occurs in the transition from uncontaminated soils to a press contamination of *I*_*C*_ = 0.0001 *mg* ⋅ *cm*^−3^ ⋅ *h*^−1^. While each 0.0001 *mg* ⋅ *cm*^−3^ ⋅ *h*^−1^ increase in *I*_*C*_ leads to further change in each of USOC, DOC, microbial biomass, extracellular enzymes and CO_2_, the relative change in each quantity becomes less with each successive jump in press contamination rate.

Lastly, the same total amount of contamination is deposited into the environment for the press versus pulse contamination scenarios, and therefore there is not much difference in the 30 year outcomes for the different quantities. But there is some difference, with a slightly better outcome for all quantities under pulse contamination. This is because, after the initial, large deposition, the soil system can begin to clear out the contamination and microbial biomass and extracellular enzymes can begin to recover and increase their amounts. The process takes thousands of years, so after 30 years, the differences are small, but differentiable. This is true for both barren and vegetated environments, although the differences are greater in vegetated environments due to the overall greater amount of soil dynamics at play.

## Acknowledgments

This work was funded by the National Science Foundation Award Nos. DMS-1853610, CNS-2018575, CBET-2329655, CBET-1603741, and DEB-2120677.

## Data Availability Statement

The code used to generate the results in this study is available from the corresponding author upon reasonable request.

## CRediT authorship contribution statement

**Sepideh Vafaie:** Conceptualization (equal); Data curation (lead); Formal analysis (lead); Investigation (lead); Methodology (lead); Software (lead); Validation (lead); Visualization (lead); Writing – original draft (lead); Writing – review & editing (equal). **Nina M. Goodey:** Conceptualization (equal); Funding acquisition (equal); Investigation (supporting); Methodology (supporting); Writing – review & editing (equal). **Jennifer Adams Krumins:** Conceptualization (equal); Funding acquisition (equal); Investigation (supporting); Methodology (supporting); Writing – review & editing (equal). **Eric Forgoston:** Conceptualization (equal); Funding acquisition (equal); Investigation (supporting);Methodology (supporting); Project administration (lead); Supervision (lead); Writing – original draft (supporting); Writing – review & editing (equal).

